# Effect of apnea and food reward on the autonomic modulation of heart rate in bottlenose dolphins (*Tursiops truncatus*) and belugas (*Delphinapterus leucas*)

**DOI:** 10.64898/2026.06.22.733846

**Authors:** Angelo G. Torrente, Bertrand Bouchard, Marine Pery, Pablo Pezzino, Julieta Arenarez, Adrian Gonzalez, Idaira, Francesco Bonadonna, Sylvia Campagna, Andreas Fahlman, Aurelie Celerier

## Abstract

1.

**Background:** Heart rate (HR) and HR variability (HRV) reflect the autonomic modulation of cardiac activity and are key indicators of physiological state and stress in vertebrates. However, measuring these parameters in cetaceans has been historically challenging due to the difficulty of recording electrocardiograms (ECGs) underwater.

**Methods:** We developed an all-in-one ECG suction-cup to record HR during apnea, respiration and food rewards in bottlenose dolphins (*Tursiops truncatus*, N = 8) and belugas (*Delphinapterus leucas*, N = 2). Moreover, we obtain comparative measurements of ECG waveform and HR oscillation in one killer whale (*Orcinus orca*).

**Results:** Species-specific ECG patterns were observed, including biphasic T waves in all three species and bifid P waves unique to belugas. All three species exhibited oscillations in resting HR and pronounced bradycardia during short apnea (∼1 min), reflecting respiratory-linked autonomic modulation. Moreover, HR oscillations persisted throughout apnea, possibly reflecting a mechanism to cyclically improve body oxygenation during diving. Short-term food deprivation had no significant effects on bottlenose dolphins and belugas’ HR, while subsequent feeding ad libitum reduced HR (by ∼20% in dolphins, ∼40% in belugas) and increased HRV, suggesting the activation of vagal modulation.

**Conclusions:** This study establishes baseline HR and HRV parameters during breathing, apnea and food reward in three cetacean species, highlighting autonomic modulation across different behaviors and conditions and validate a non-invasive tool for studying marine mammal physiology.

## Introduction

Cetaceans have evolved remarkable physiological adaptations to sustain a fully aquatic lifestyle while remaining air-breathing mammals (Fahlman, 2024). These adaptations include intermittent or apneustic respiration and multiple mechanisms that optimize oxygen consumption, such as the expression of specific hemoglobin isoforms and a tight control of heart rate (HR) and metabolism (Williams and Davis, 2021). Among these traits, the regulation of HR has long been recognized as central to diving capacity in marine mammals (Scholander, 1940; Irving et *al*., 1941; Ponganis, 2011), having been described as the “master switch of life” (Scholander, 1963).

As in other mammals, HR regulation in cetaceans is primarily driven by autonomic modulation of the heart’s pacemaker activity (Aoki et al., 2021; Bickett et al., 2019; Fahlman et al., 2020). Indeed, the autonomic nervous system, a component of the peripheral nervous system, regulates involuntary physiological processes such as HR, blood pressure, respiration, digestion, and sexual arousal (Waxenbaum et al., 2025). Autonomic output varies dynamically with the body’s need to respond to internal and external stimuli. Therefore, the analysis of HR variability (HRV) provides a powerful, non-invasive window into autonomic modulation (McCraty and Shaffer, 2015). In terrestrial mammals a decrease of HR and an increase of HRV are generally considered a consequence of a raise in vagal tone (parasympathetic), while the increase of HR and the decrease of HRV are generally related to the predominance of adrenergic (sympathetic) activity (McCraty and Shaffer, 2015). However, these correlations have been studied only in terrestrial animals that maintain a continuous and cyclical respiration, and could thus not be immediately applicable to the autonomic modulation during apnea (Cunha Oliveira et al., 2026).

HRV is also widely recognized as a marker of cardiovascular health, with higher HRV generally indicating better cardiac adaptability and lower HRV being associated with stress, impaired autonomic regulation and aging (McCraty and Shaffer, 2015). This relationship has been demonstrated in humans and other mammals, including dogs, and rodents (Haensel et al., 2008; Pizzo et al., 2022; Umesh et al., 2025).

In continuous breathers (terrestrial mammals), autonomic control of HR at rest is closely linked to respiration through the physiological mechanism of respiratory sinus arrhythmia (RSA) (Cirus Kauveh Shiran, B. S., 2025). RSA produces a brief rise in HR during inspiration followed by a comparable decrease during expiration, a process thought to improve ventilation–perfusion matching and enhance gas-exchange efficiency (Ben-Tal *et al*., 2012). Similar respiration-linked HR fluctuations, analogous to but not necessarily identical to classical respiratory sinus arrhythmia (RSA), have been described in cetaceans and are characterized by a rapid HR acceleration with each breath, followed by a gradual decline until the next breath (Fahlman et al., 2023), a mechanism also known as “respiratory tachycardia” (Burggren et al., 2024). Compared to terrestrial mammals, fluctuations in HR at each cycle of respiration are particularly pronounced, especially when the interval between two breaths includes a dive. However, only a few studies have explicitly analyzed these patterns.

Intrigued by this exceptional HR modulation in cetaceans, we investigated their autonomic regulation during respiration and apnea, and using food as a strong rewarding stimulus that could activate the autonomic nervous system. To achieve this, we developed a new all-in-one suction-cup capable of recording high-quality electrocardiograms (ECGs) underwater. Using this device, we measured HR and the diving response in a group of eight bottlenose dolphins, two belugas, and one killer whale maintained under managed care. In bottlenose dolphins and belugas, we further examined the autonomic response to food reward by analyzing HR and HRV dynamics.

## 2. Materials and methods

### (a) Animals used in the study

We studied a group of eight bottlenose dolphins (*Tursiops truncatus*, all males) and two belugas (*Delphinapterus leucas*; one male and one female) housed at the Oceanogràfic of Valencia (Spain). In a second phase of the study, we included also one female killer whale (Orcinus orca) housed at Loro Parque (Tenerife, Spain), to had a third species of comparison for the ECG waveform and the fluctuation of HR during respiration and apnea. Before beginning the ECG recordings, all study protocols were reviewed and approved by the Animal Care and Welfare Committee of the Oceanogràfic (protocols OCE-13-23) and by the French National Ethical Committee (Comité Consultatif National d’Éthique, permit No. 1286.5392). All procedures, animal husbandry, and management practices were discussed and pre-approved by the attending veterinarians and animal care teams at both facilities, who carefully supervised the study. Five of the dolphins and one beluga were born under human care. Three dolphins and one beluga were wild-born and have lived under professional care in aquariums for over 20 years. The killer whale was rescued from the wild after being found in distress. All animals were fed ad libitum a mixed diet (fish, squid, octopus, and gelatin) several times per day.

### (b) All-in-one suction cup for ECG recordings

Most previous studies on cetacean cardiac physiology used custom-built ECG systems consisting of two suction-cup electrodes connected by cables to an external recording device attached to the animal’s skin, typically on the sternum and between the pectoral fins (Bickett et al., 2019; Blawas et al., 2021; Elmegaard et al., 2019, 2021; Lyamin et al., 2016; Williams et al., 2015). Although successful in some cases, this configuration can introduce electrical noise caused by cable movement between the electrodes and the recording unit and can expose the interconnection points to seawater, increasing the risk of short circuits and signal loss.

To obtain high-quality ECG recordings in the three species of cetaceans analyzed we developed an elliptical all-in-one suction cup designed to house a waterproof chamber containing an archival voltage-difference recorder for ECG measurements. We molded the suction cup from silicone (Reschimica Srl; Barberino Tavarnelle, Florence, Italy) in our laboratory, using custom made templates. The elliptical suction cup measured approximately 18 × 11 cm at the base (suction rim) and on the ventral surface, presented two helicoidal stainless-steel spring electrodes that allowed us to record ECGs from the contact with the animal’s skin. By integrating all electronic components within a single waterproof suction cup, we minimized cable movement and exposure of connectors to water, thereby optimizing the signal quality and reliability of ECG recordings. In addition, the device’s easy attachment and removal made the system completely non-invasive. After trying different positions, we found that having the electrodes placed on the longitudinal axis of the body in between the pectoral fins, was the best way to obtain high-quality ECGs (**Fig. 1**). Under these settings we recorded the HR of eight bottlenose dolphins, two belugas and one killer whale (**Fig. 1**).

**Fig. 1:**
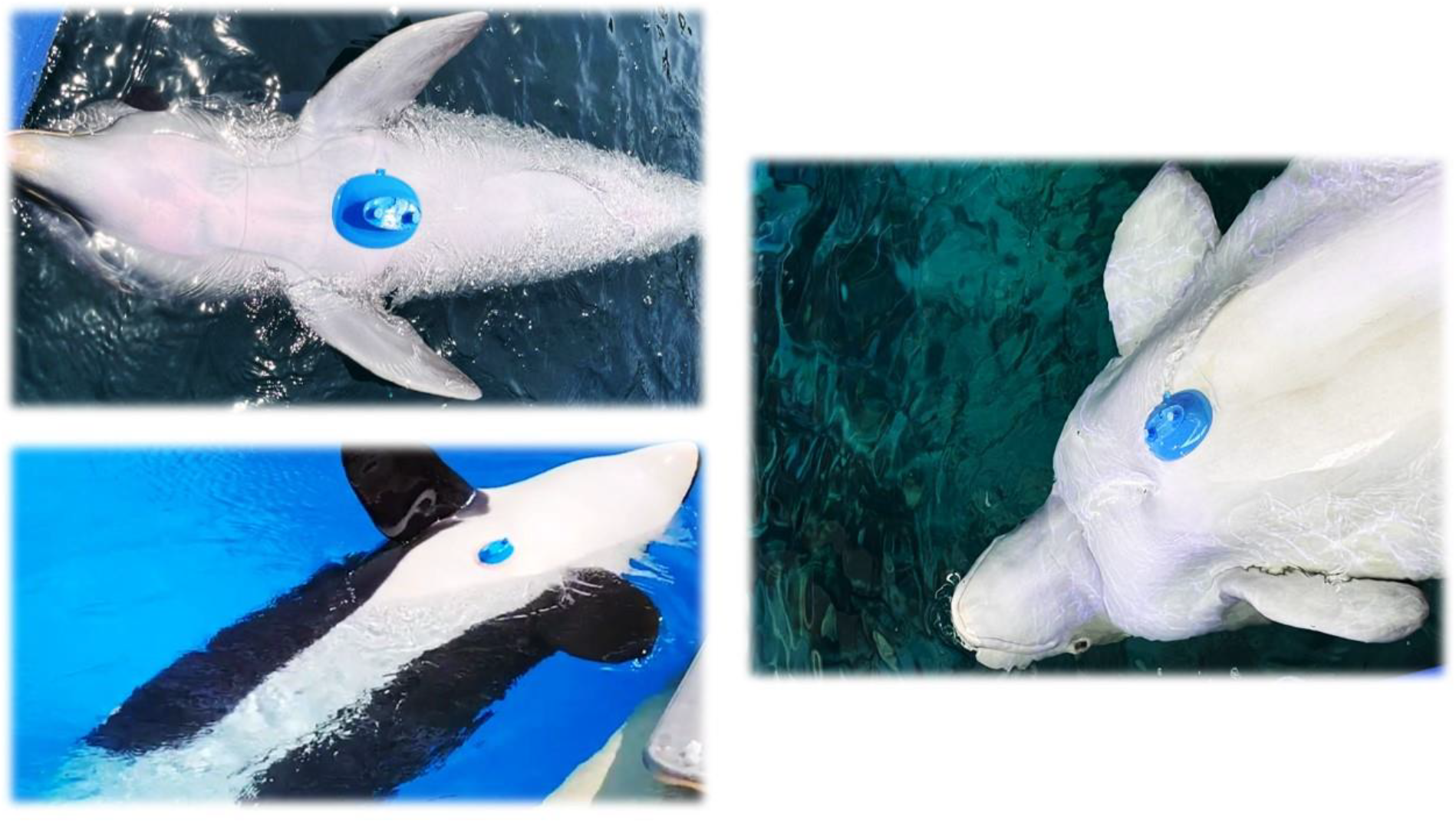
ECG device placement. Prototypes of the ECG suction cup, placed on the thoracic area of bottlenose dolphins (*Tursiops truncatus*), belugas (*Delphinapterus leucas*) and killer whales (*Orcinus orca*)

As in our previous work (Bonadonna et al., 2024), we obtained ECG signals using the commercially available miniwaves EMG device (Cometa Srl; Milan, Italy). Although originally designed for electromyography, this miniaturized and waterproof logger provides high-quality ECG traces when positioned near the thoracic region. The device recorded voltage differences at a fixed sampling rate of 2000 Hz and included a triaxial accelerometer (sampling rate = 142.8 Hz; range = ± 8 g). The logger could transmit ECG and accelerometer data via Wi-Fi to a receiver connected to a laptop when used out of the water, or operate as an autonomous archival unit using its internal memory, as in the present experiments.

### (c) Experimental protocols

The ECG suction cup was placed on the animal’s sternal area, between the pectoral fins. Each session was generally completed within 15 minutes, including having the animal in a ventral posture, attaching the suction cup, and recording the ECG. Animals were not restrained and could interrupt the session at any time without any negative consequence. Trainers continuously monitored the animals and interrupted the session if any sign of distress was detected. Before the experiments, all bottlenose dolphins and belugas had been desensitized to the contact with the ECG suction cup through repeated training sessions during the previous year. The killer whale had already participated in other studies involving suction cup attachments to the ventral area, and was therefore considered desensitized to this procedure.

During each trial, the selected animal remained stationary and floating near the edge of the pool and in contact with the trainer. Other animals present in the facility were kept under trainers’ control, either in distant corners of the same pool or in adjacent pools, in order to allow vocal contact with the focal animal, while minimizing disturbance during the test. Each trial began with an initial apnea of at least one minute, with the animal’s ventral side oriented upward and partially emerged (“ventral position”). During this period, we attached the suction cup to the thoracic area, and we recorded a reference HR under apnea. The animal was then asked to rotate to “dorsal position”, exposing its dorsal side above the water, and was maintained at the surface with its blowhole emerged to allow breathing at their voluntary frequency.

#### Experiment 1. Comparison of apnea and breathing

The influence of apnea on the autonomic modulation of heart activity was studied during surface rest by comparing two different conditions: “**Apnea**” in ventral position with the blowhole underwater and “**Breathing**” when animals were then placed in dorsal position in order to be able to breathe at their voluntary frequency. In both conditions, animals were fed ad libitum as during usual training sessions.

#### Experiment 2. Effects of food reward

In this experiment we used food reward to study the autonomic response to external stimuli. This was done by measuring HR and HRV under two trial conditions: a short food deprivation phase (**No Food**) and a period of ad libitum food reward (**Food Reward**). The “**No Food**” condition generally lasted 2 minutes, while “**Food Reward**” lasted at about 1 minute. During this time the animals were maintained in dorsal position with no other external stimuli from the trainers (no speaking, or petting), except for the minimal physical contact required to keep the animal stationary.

### (d) ECG traces

ECG traces represent the temporal variation of the electrical potential difference between two contact points on the skin, typically located on the thorax or at body extremities (arms and legs). In our experiment, these two points corresponded to the stainless-steel spring electrodes positioned at the two extremities of the elliptical suction cup, which were in direct contact with the cetaceans’ skin. Data collected by the Miniwave recorder were downloaded using the dedicated software EMG and Motion Tools (Cometa Srl; Milan, Italy). As in humans and other animals, we were able to identify main deflections across the ECG trace in both positive and negative directions, corresponding to the ventricular depolarization that triggers ventricular contraction. These peaks are typically used to mark each heartbeat and are referred to as “R waves”.

In addition, in all recordings obtained during apnea (device out of the water) and in most recordings collected with the device immersed, we were able to distinguish smaller waves that preceded the R wave and displayed a longer decay. These “P waves” correspond to the atrial depolarization, which induces atrial contraction. Following the R wave, we also detected longer-decay deflections commonly referred to as “T waves”, corresponding to ventricular repolarization and relaxation. R waves were automatically detected using the ECG Analysis or HRV modules of LabChart Pro 8 (ADInstruments Ltd; Dunedin, New Zealand). When signal noise reduced the accuracy of automatic detection, R waves were manually added or excluded using the HRV module of the same software.

### (e) Statistics and data analysis

We reported all data as Mean ± Standard error of the mean (SEM). Statistical tests, such as t-test were applied as indicated in the text and figure legends and were performed using GraphPad Prism 10 (GraphPad Software, Boston, Massachusetts, USA). The capital letter “N” indicates the number of animals included in each test, while the lowercase “n” indicates the number of trials or repetitions performed. Following best practices for HR measurements (Chabot et al., 1991), we calculated the average HR under all experimental conditions by dividing 60 by the mean RR interval expressed in seconds. For these analyses, representative 1-min segments were randomly selected from artifact-free periods of the ECG recordings. Global HRV was calculated as Standard Deviation of RR interval (StDRR) or better as the coefficient of variability of RR intervals (CVRR) = StDRR/meanRR*100, which represents StDRR normalized by the Mean RR.

Beat-to-beat HRV was calculated using the parameter RMSSD 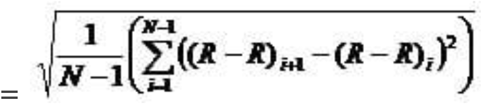. To avoid the influence of the RR mean on RMMSD when comparing condition that have a strong difference of HR, such as **Apnea** and **Breathing**, we also included the RMSSD normalized by the average RR (RMSSD/Mean RR) to confirm that the significant changes of RMSSD were real and due to a strong change of Mean RR.

Simultaneous accelerometer data were used to assess the animal’s posture and when possible to determine the timing of respiratory events.

## 3. Results

### (a) ECG profiles in three cetacean species

We started by analyzing ECG recordings from bottlenose dolphins and belugas under the condition of “**Apnea**” and “**Breathing**” described in the **Experiment 1** (see methods) to identify potential differences with the ECG waveforms typically observed in humans. The examination of the ECG waveform revealed the main phases of cardiac electrical activity: P wave (atrial depolarization), R wave (ventricular depolarization), and T wave (ventricular repolarization), already known in human and other vertebrates (**Fig. 2A**) (Bickett et al., 2019; Bonadonna et al., 2024). Consistent with previous findings (Bickett et al., 2019), the ECG signal amplitude was higher when recordings were obtained in the ventral position (**Apnea**) and the ECG suction cup was exposed above the water, than in the dorsal position (**Breathing**; **Fig. 2A**). However, the overall waveform morphology remained similar under both conditions. Both species exhibited an inverted or biphasic T wave (**Fig. 2A**), a feature that we confirmed also in one killer whale. This kind of waveform for the ventricular repolarization have been already described in early bottlenose dolphins’ ECGs (Ridgway, 1968) and in human it is often associated with myocardial ischemia or hypokalemia in humans (Kotsialou et al., 2024). However, because we consistently observed this inverted or biphasic of T wave in several individuals, supposedly in good health, of three cetacean’s species, it could be a species-specific feature for cetaceans. Moreover, belugas showed a distinctive double (bifid) P wave, which could be caused by the large atrial size, which could induce asynchronous, but not necessarily pathological, conduction of the pacemaker impulse from the sinoatrial node to the right and left atria.

**Fig. 2:**
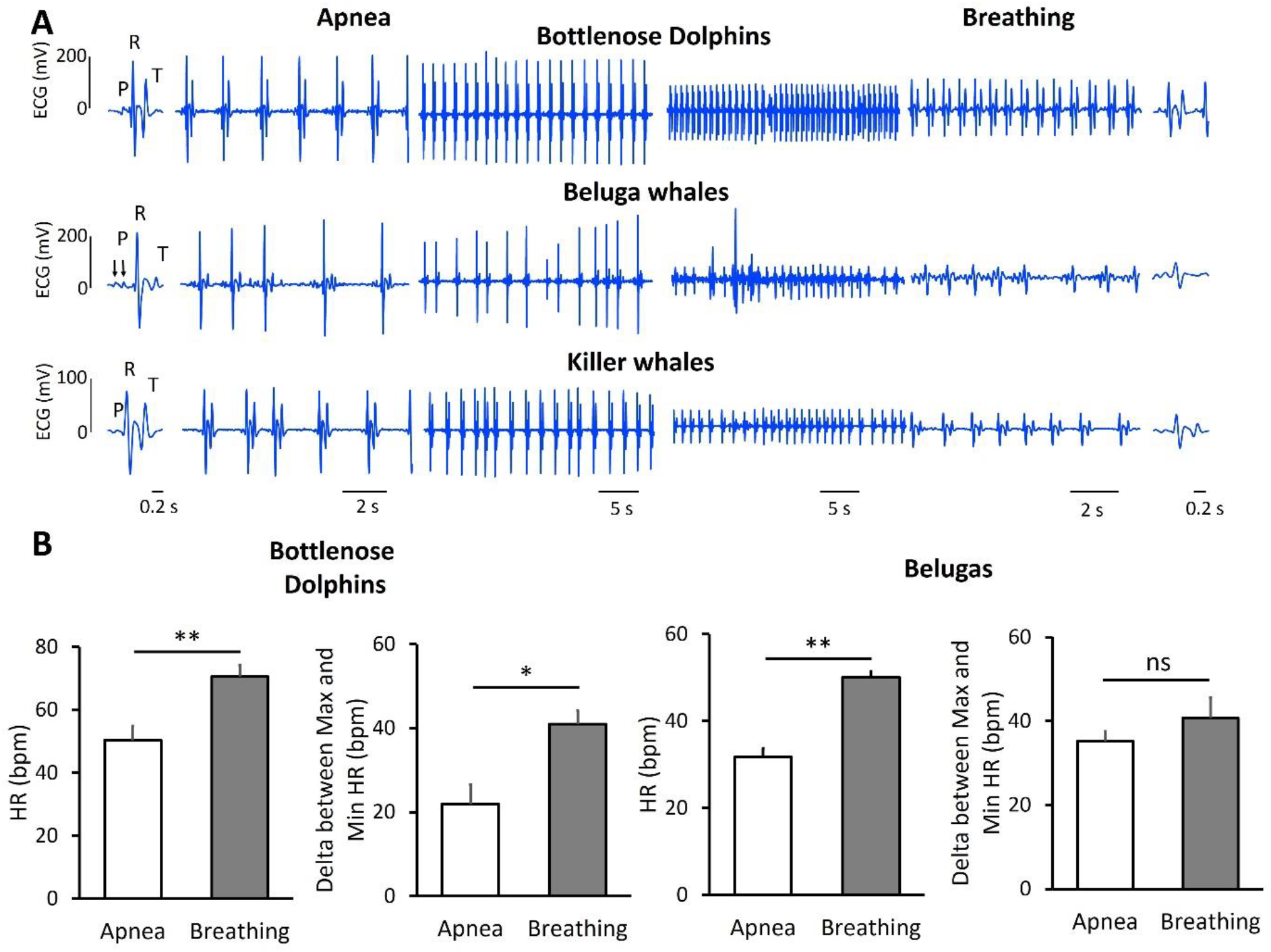
**A**, Representative ECG traces recorded from bottlenose dolphins, belugas and killer whales resting at surface, under breathing or apnea, and at different time scales. Biphasic T waves were observed in all three species, whereas double (bifid) P waves were present only in belugas (black arrows). **B**. Comparison of HR and the delta between maximal (Max) and minimal (Min) HR under Apnea and Breathing in bottlenose dolphins and belugas. Bottlenose dolphins, N = 8, belugas, N = 2, n = 5; **p<0.01 and *p<0.05 by paired t-test.

### (b) Experiment 1: Comparison of apnea and breathing

When comparing apnea and breathing conditions in bottlenose dolphins and belugas, as expected we observed a lower HR under apnea (**Fig. 2**). Moreover, we observed HR fluctuation that we initially characterized by comparing maximal and minimal HR for each recording. The delta between maximal and minimal HR was significantly more pronounced during breathing than apnea in bottlenose dolphins, but was similar during apnea and breathing when looking to belugas (**Figs. 2B**).

While HR fluctuations during breathing could be attributed to the cyclical changes associated with respiration, it was unexpected to observe a comparable degree of variability during apnea **(Fig. 3)**. In particular we observed that about half of the dolphins recorded maintained clear cyclical HR variations even under apnea with a frequency of 8.3 ± 0.4 cycles/min N=5, n=7 (**Fig. 3**), while in the other dolphins we observed only little changes of HR. In belugas we also observed alternation between slow and fast HR at each cycle of respiration (**Fig. 3**). Moreover, during apnea, belugas showed abrupt changes of HR consistent with single, doublets or triplets of closely spaced heartbeats, repeated at 6.6 ± 0.7 cycles/min N=2, n=8 (**Fig. 3**) with similar instantaneous HRs. This alternation between slow and fast HR caused fluctuation between maximal and minimal HR similar to what recorded during voluntary respiration although with a faster cycle (**Fig. 2B and 3**). We confirmed a similar pattern of HR fluctuation during apnea also in one killer whale.

**Fig. 3:**
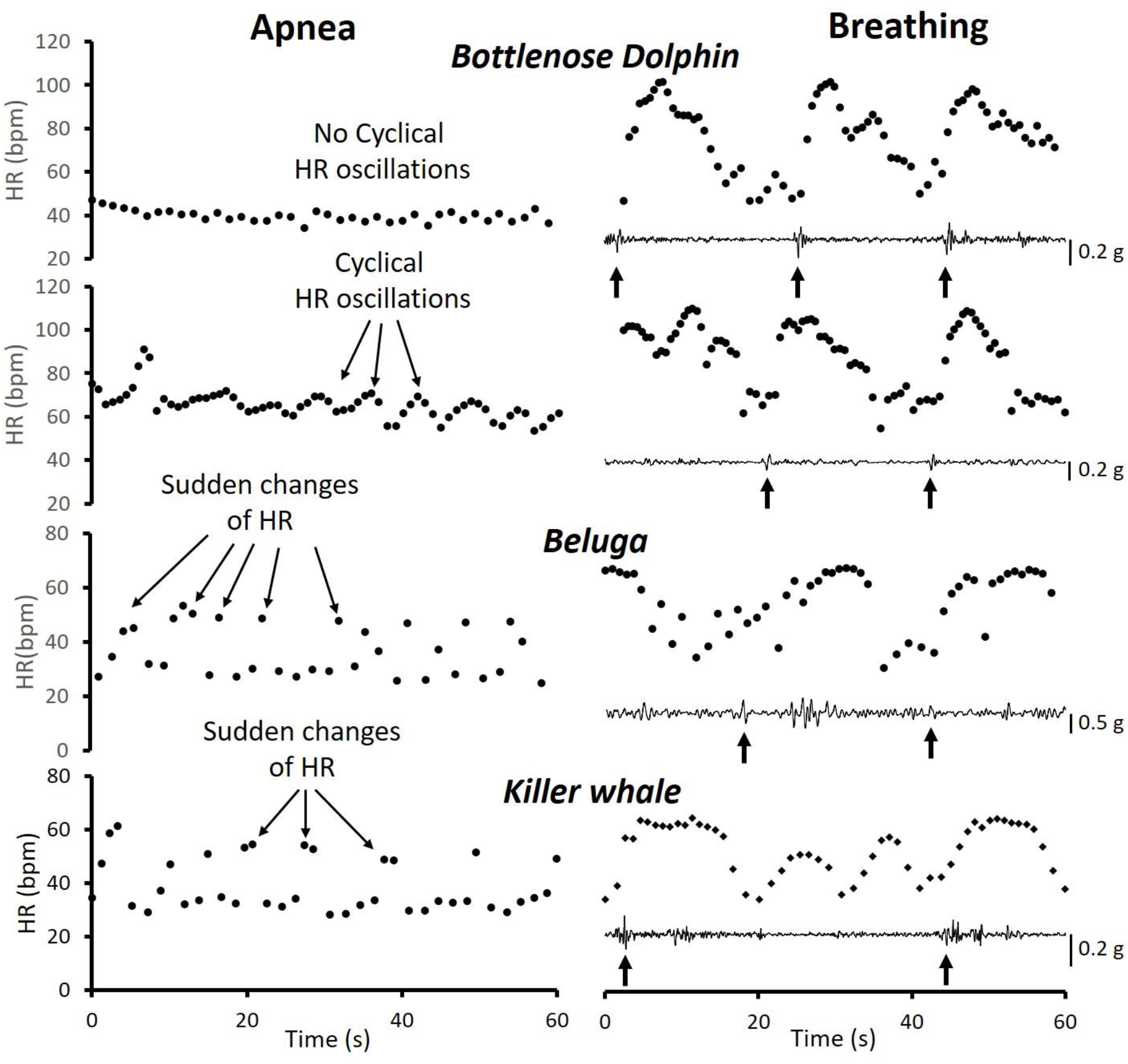
Instantaneous HR recorded in three cetacean species in apnea and breathing conditions. In the right-hand panels, acceleration traces are shown below the HR plots; black arrows mark the likely timing of respiratory events.

Accordingly, to the outcome obtained measuring the difference between maximal and minimal HR, when comparing breathing and apnea we observed a decrease of global HRV only in bottlenose dolphins, when using the Coefficient of Variability (CVRR) that normalize the Standard deviation of RR intervals (StDRR) to the Mean RR, as we already reported (Cunha Oliveira et al., 2026). While we observed no significant changes of HRV in belugas (**Fig. 4**). Whether in bottlenose dolphins, belugas or the killer whale, the fluctuation of HR during apnea had a cyclical period faster than the typical period between two respirations (**Fig. 3**).

**Fig. 4:**
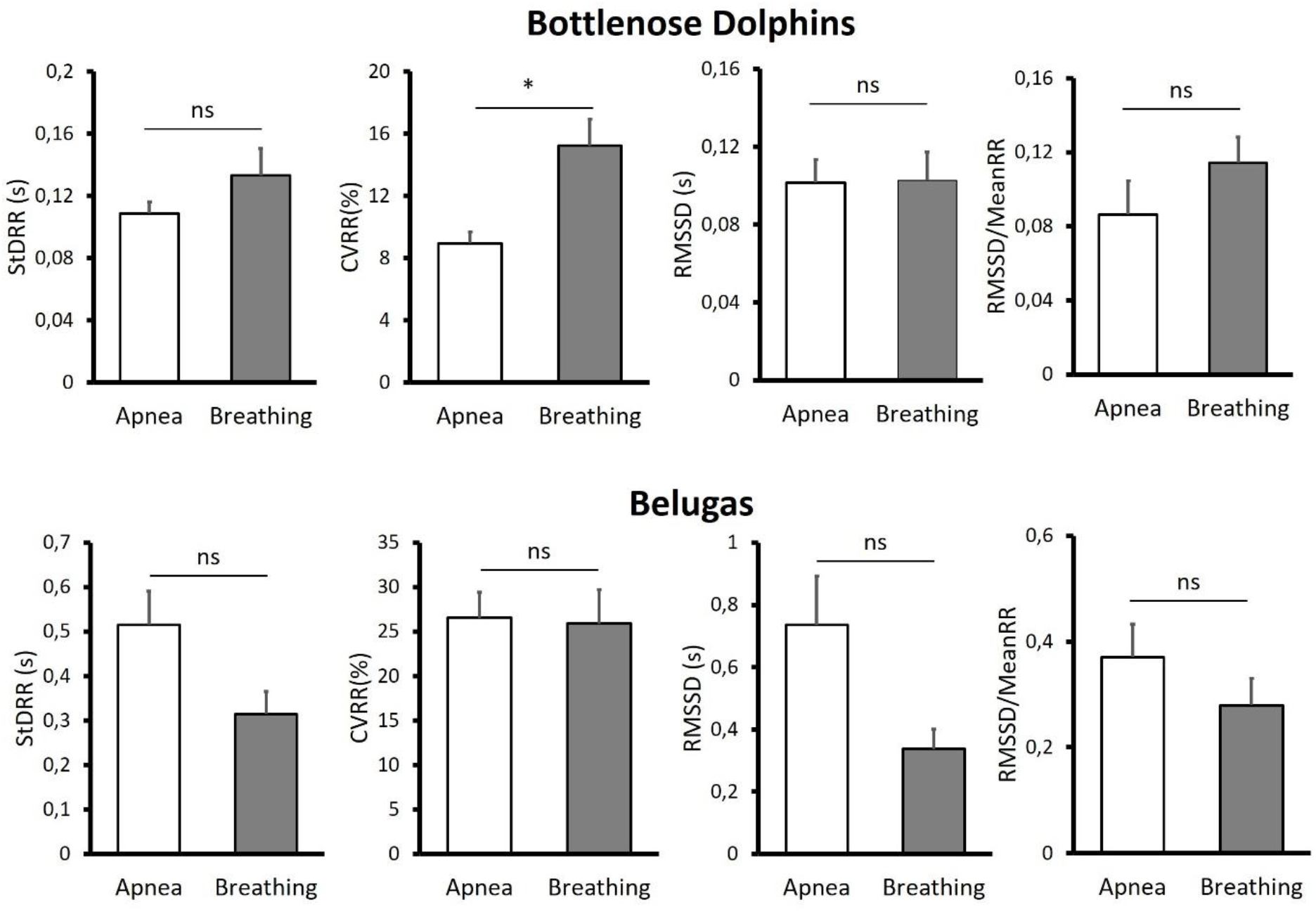
Parameters of global (StDRR and CVRR) and beat-to-beat HRV (RMSSD and RMSSD/MeanRR) in Bottlenose dolphins and belugas during apnea and breathing conditions of experiment 1. Bottlenose dolphins, N = 8, belugas, N = 2, n = 5; *p<0.05 by paired t-test

### (c) Experiment 2: Effects of food rewarding

We then further investigated the modulation of cardiac activity by the autonomic nervous system by comparing HR during a short phase of no feeding with a subsequent phase of feeding ad libitum, used as a strong reward stimulus. During the whole protocol of **Experiment 2** (see methods) the animals were maintained resting in dorsal position and were free to breathe at their voluntary frequency. Under the “**No food**” condition in both species, HR did not differ significantly from those observed in the “**Breathing**” condition described in Experiment 1 (in Bottlenose dolphins: 80 ± 3 vs. 71 ± 4 bpm, N = 8; p=0.13 by paired t-test. In belugas: 54±1 vs. 51±1 bpm, N=2, n=3; p=0.15 by paired t-test; **Fig. 5**).

**Fig. 5:**
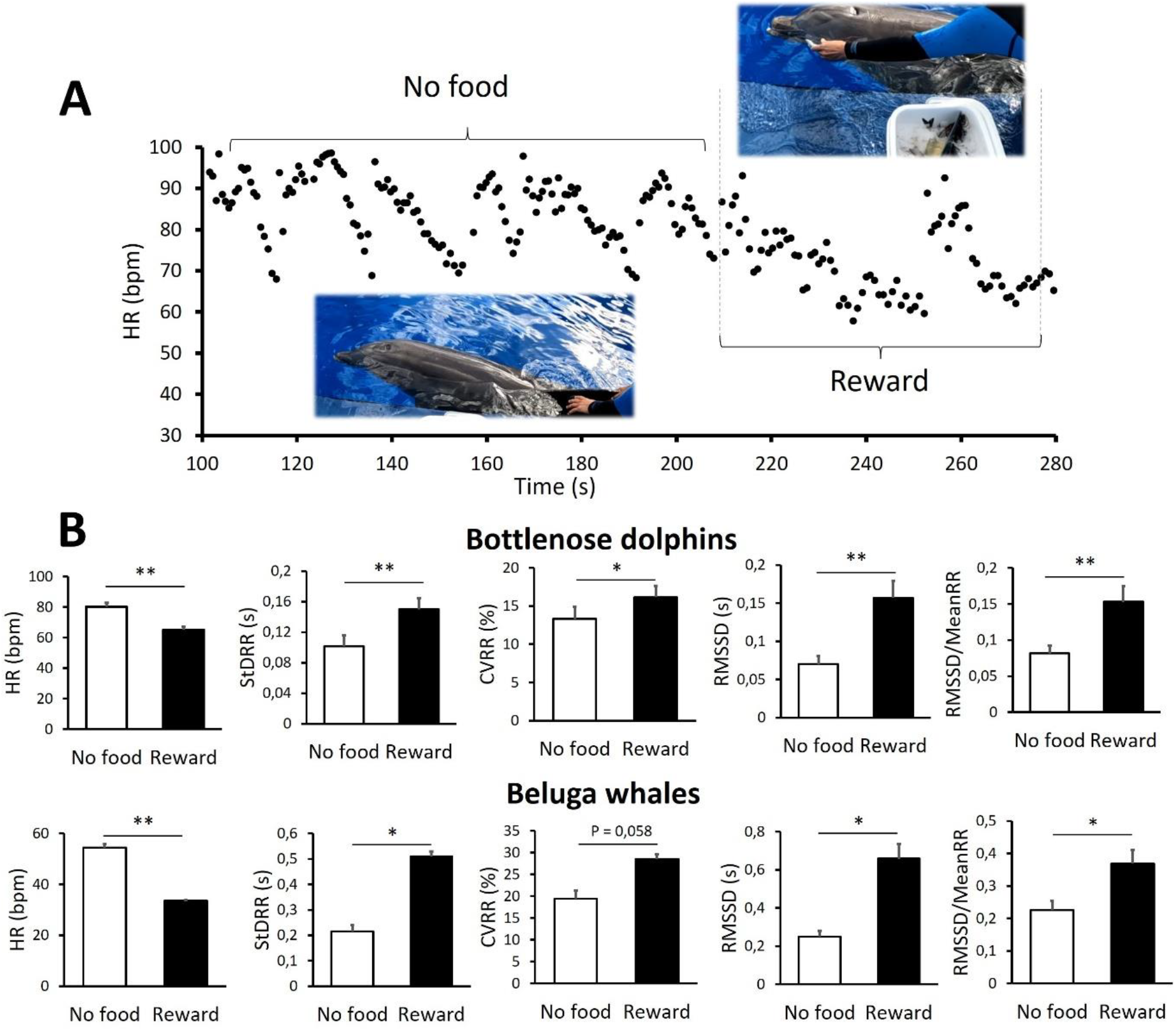
A Instantaneous HR over time in an experimental protocol comparing conditions before and after food reward in bottlenose dolphins. **B**. HR and HRV parameters (StDRR, CVRR, RMSSD and RMSSD/MeanRR) measured before (No food) and after food reward (Reward) in bottlenose dolphins and belugas. **p<0.01 and *p<0.05 by paired t-test. ns = non-significant. N = 8 for Bottlenose dolphins and N = 2, n = 3 for belugas.

In Bottlenose dolphins, the transition from No food to Food reward caused an immediate decrease in HR and a simultaneous increase in HRV (**Fig. 5A**). Specifically, the parameters StDRR and CVRR, reflecting global HRV, and RMSSD and RMSSD/MeanRR, representing beat-to-beat variability, all increased during feeding (**Fig. 5B**), contrary to what we observed when comparing breathing with apnea. The decrease of HR and the specific increase of RMSSD, which in terrestrial mammals is primarily associated with parasympathetic (vagal) activation, suggest that feeding shifted the autonomic balance toward enhanced parasympathetic influence. Similar, food reward in belugas induced HR decrease and HRV increase (**Fig. 5B**).

Interestingly, in both species, food reward also produced a significant decrease in both Min and Max HR, suggesting that the autonomic activation elicited by feeding induced a general shift of the HR pattern toward lower values (**Fig. 6**).

**Fig. 6:**
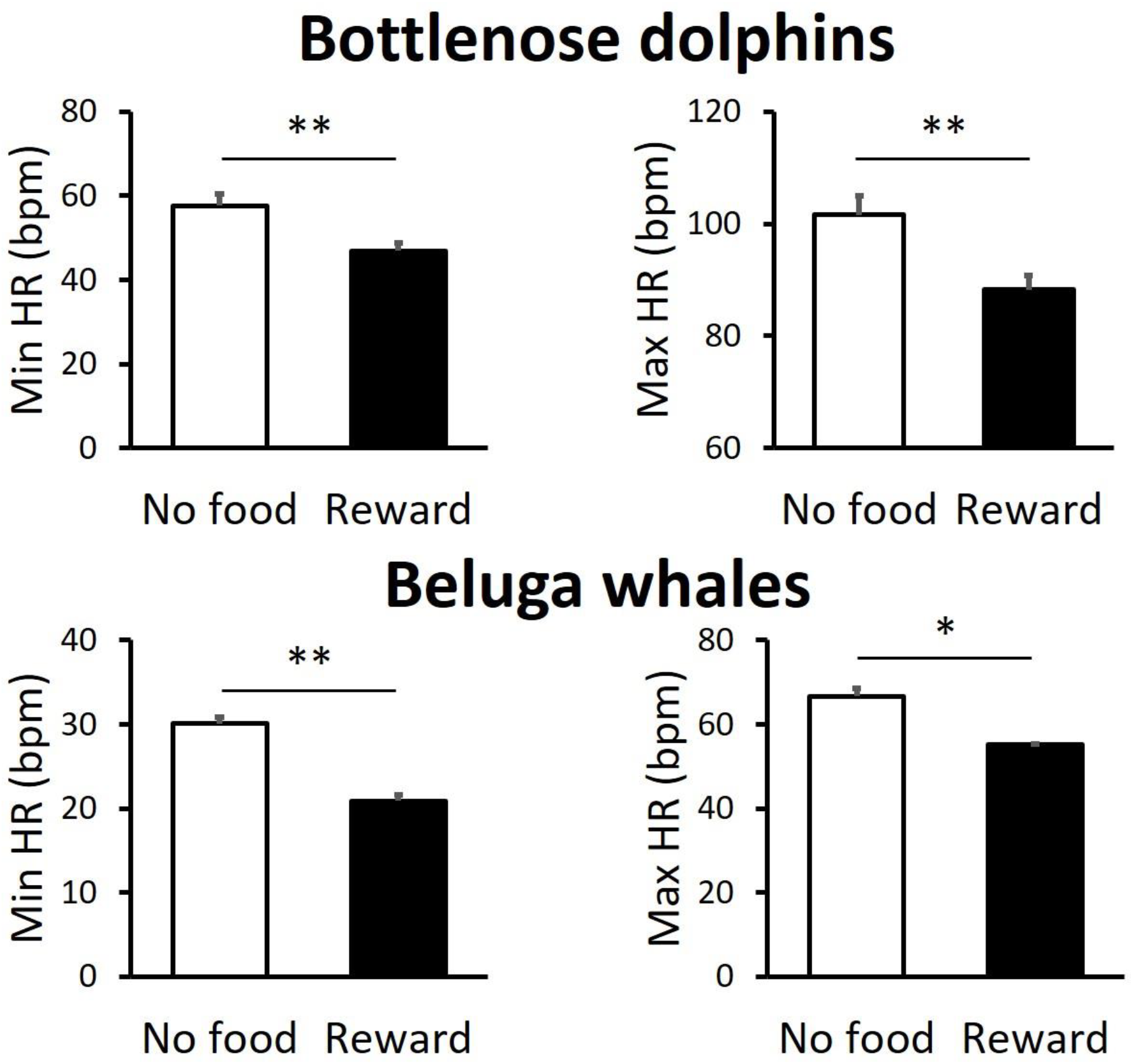
Minimum and maximum HR measured before (No food) and after food reward (Reward) in bottlenose dolphins and belugas. **p<0.01 and *p<0.05 by paired t-test. N = 8 for bottlenose dolphins and N = 2, n = 5 for belugas.

## 4. Discussion

Understanding how cardiac rhythm is modulated in marine mammals provides crucial insight into the physiological mechanisms that enable these animals to sustain prolonged dives, optimize oxygen use, and respond to environmental stimuli. Using an innovative, non-invasive ECG recording system, we analyzed HR and HRV in bottlenose dolphins (*Tursiops truncatus*) and belugas (*Delphinapterus leucas*). The compact and waterproof design of the elliptical suction cup developed for this study enabled high-quality ECG recordings with minimal electrical interference or discomfort for the animals.

The ECG analyses revealed cetacean-specific cardiac waveforms, including a biphasic and/or inverted T wave consistently observed in bottlenose dolphins (*Tursiops truncatus*) and belugas (*Delphinapterus leucas*), but also in one killer whale. Although this morphology is often associated with myocardial ischemia or electrolyte imbalance in humans (Kotsialou et al., 2024), similar T-wave patterns have previously been reported in healthy Bottlenose dolphins (Ridgway, 1968) and pinnipeds such as the walrus (*Odobenus rosmarus*) (Storlund et al., 2021), as well as in terrestrial mammals including dromedary camels (*Camelus dromedarius*) (Al Khamis et al., 2023) and horses (*Equus ferus caballus*) (Nissen et al., 2022). This morphology therefore likely represents a species-specific electrophysiological adaptation rather than a pathological condition. It may arise from unique anatomical and functional characteristics of the cetacean heart, such as a size, high ventricular mass, complex ventricular geometry, or specialized Purkinje fiber distribution. In addition, the predominant parasympathetic tone characteristic of diving mammals (Ponganis et al., 2017) could also slowdown ventricular repolarization and contribute to this distinctive T-wave profile compared to humans.

Belugas also exhibited a double (bifid) P wave, suggestive of asynchronous depolarization of the right and left atria, potentially linked to their large atrial size and/or specialized conduction pathways between the sinoatrial node and the atria (Meek, 2002). Because all studied animals were in confirmed good health, these features likely represent normal interspecific variation rather than pathology. Though, future work including additional individuals will be necessary to confirm these patterns.

HR measured at the surface for bottlenose dolphins (*Tursiops truncatus*) and belugas (*Delphinapterus leucas*) was comparable to values previously reported in the literature (Blawas et al., 2021b; Fahlman et al., 2020) although higher than others (Bickett et al., 2019). As previously noted (Aoki et al., 2021), these HRs exceeded those predicted for terrestrial mammals of similar body mass based on the allometric equation 241 × [body mass (kg)]^−0.25^(Aoki et al., 2021; Stahl, 1967). Based on this relationship, the expected resting HR would be approximately 64 bpm for a 200-kg bottlenose dolphin, 40 bpm for a 1300-kg beluga (average female), and 32 bpm for a 3000-kg killer whale (average female). Interestingly, in bottlenose dolphins and belugas, the predicted values were intermediate between the HRs observed during apnea and those recorded during breathing (**Fig. 7**). While the value recorded in a single killer whale were slightly higher of the expected allometric estimation.

**Fig. 7:**
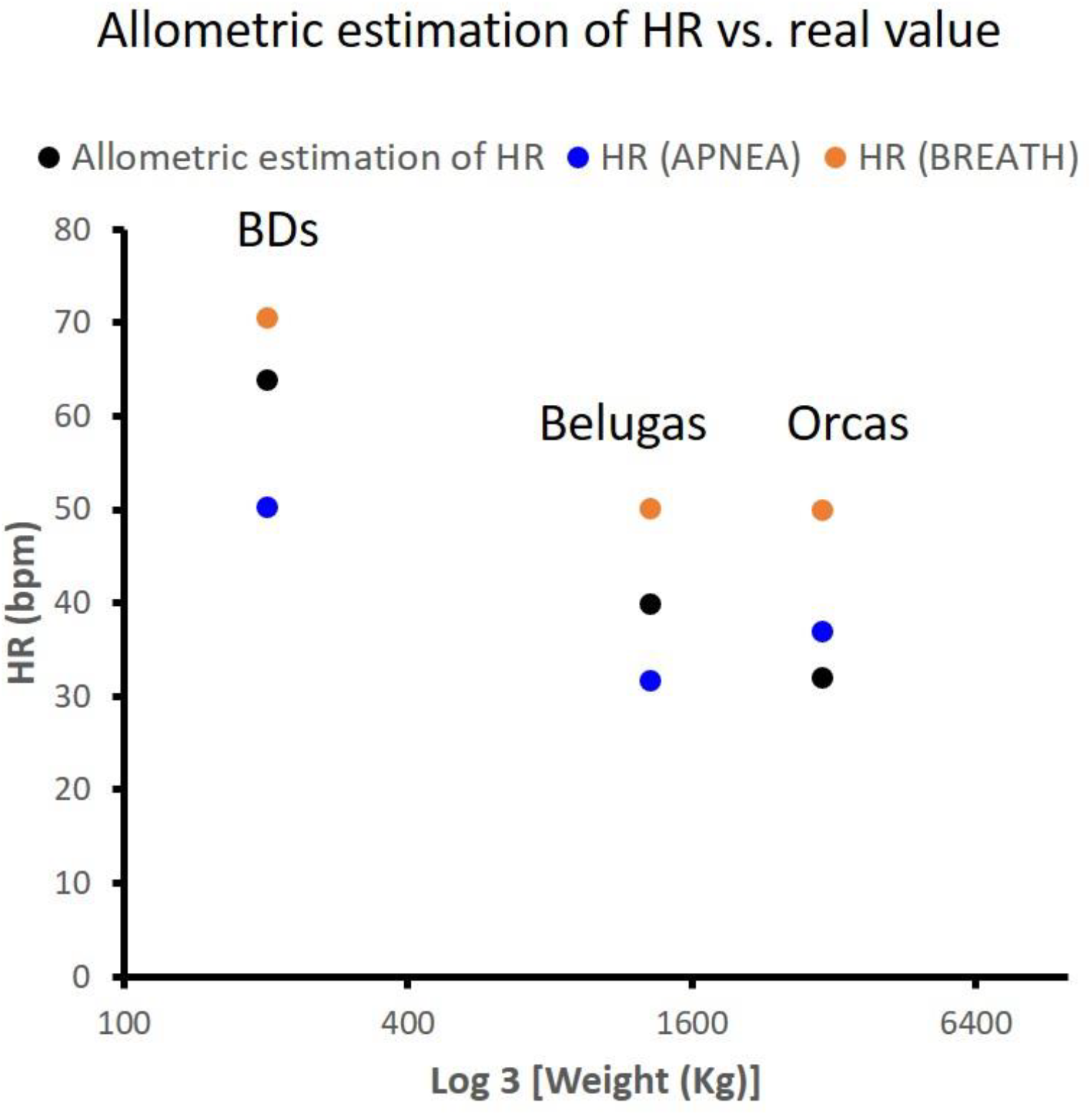
Allometric estimation of HR: Comparison between HR estimated by the Stahl allometric equation 241 × [body mass (kg)]^−0.25^, versus real value recorded in Bottlenose dolphins, belugas and killer whale.

In bottlenose dolphins (*Tursiops truncatus*), belugas (*Delphinapterus leucas*) and one killer whale, HR oscillations were associated with respiration and the mechanism of respiratory sinus arrhythmia, consistent with previous studies (Blawas et al., 2021b; Fahlman et al., 2020, 2023). However, we unexpectedly observed oscillations or distinctive alternation between slow and fast HRs, even under apnea. Similar oscillations have previously been reported in cetaceans and marine mammals (Bickett et al., 2019; Blawas et al., 2021a; Storlund et al., 2026). Although their origin remains uncertain, they could result from cyclical compensation between arterial pressure and sinoatrial activity (Blawas et al., 2021a), consistent with the Mayer waves described in humans, rabbits, and rats (Julien, 2006). Further investigations will be required to test this hypothesis and clarify the physiological role of this pattern. But, because, at least in belugas and killer whales, the faster heartbeats were repeatedly interspersed within periods of pronounced bradycardia, they may help sustain oxygenation of bloodstream during apnea. The sudden alternation of fast and slow heartbeats has been previously explained as a conflict between sympathetic and parasympathetic modulation (Blawas et al., 2021a). Such a conflict or the cyclical change of vagal tone could explain the gradual change that we observed in bottlenose dolphins during apnea. However, it will hardly allow the abrupt changes observed in belugas and one killer whale. Thus, given the very fast alternation between heartbeats of about 20 bpm of difference, a particularly intriguing hypothesis is that these abrupt changes result from a quick shift in the dominant sinoatrial pacemaker region triggering heartbeats (Baudot et al., 2020; Torrente et al., 2015). Indeed, during apnea the repressive modulation of cardiac pacemaker activity mediated by vagal tone, could induce competition between different leading regions of the sinoatrial node, giving rise to sudden changes of HR with two dominant frequencies of instantaneous HR. Specifically, the slower and more represented frequencies of heartbeats would correspond to the pacemaker slowdown induced by the increase of vagal tone, while the sudden increase of HR would be generated by a group of SAN cell that would be intermittently escape the parasympathetic modulation of pacemaker activity, perhaps in correspondence of short decrease of vagal tone.

Finally, we used food reward as a simple and biologically relevant stimulus to investigate autonomic modulation in bottlenose dolphins and belugas. Compared with a short period of no food while resting at surface, food reward induced a marked decrease in HR together with an increase in HRV, a pattern consistent with enhanced parasympathetic (vagal) modulation different from what observed during apnea. These findings indicate that HR and HRV are sensitive markers of autonomic modulation in cetaceans and may provide important tools to evaluate cardiovascular dynamics related to oxygen availability or physiological responses to external stimuli. More broadly, HRV analysis could offer new opportunities to investigate, stress, motivation, and welfare in marine mammals under both managed-care and natural conditions.

## 5. Limitations

These findings should be interpreted cautiously given the limited sample size, particularly for belugas and the single killer whale.

In particular, results obtained in belugas could derive from the parental relation (mother and calf) between the two individuals included in the study. Therefore, the bifid P wave observed in both individuals may either represent a species-specific characteristic or reflect a congenital anomaly. Further recordings from unrelated belugas will be required to determine whether this ECG feature is representative of the species as a whole.

Another limitation concerns the interpretation of the cardiac responses observed during food reward. As mentioned above, except during the apnea trials, animals were free to breathe at their voluntary rate under all conditions. During food reward (ad libitum feeding) both bottlenose dolphins and belugas voluntarily reduced their breathing frequency. Because cetaceans can feed while holding their breath underwater and because the respiratory and digestive tracts are anatomically separated feeding itself does not mechanically restrict ventilation. Instead, this reduction in breathing frequency likely reflected a behavioral choice, with the animals directing their attention toward feeding rather than respiration.

This reduction in breathing frequency may have partly contributed to the HR decrease observed during feeding. However, the different dynamics of HRV indices when comparing breathing with apnea or no food with food reward indicate that food reward influenced HR and HRV likely by enhancing vagal tone as suggested by the overall decrease in both maximum and minimum HR and RMSSD increase.

## 6. Future directions

Data not shown indicated the feasibility to record HR in free swimming cetaceans using our all-in-one ECG suction-cup, including in large mysticetes. Coupling HR and HRV measurements with accelerometry, acoustic monitoring, behavioural observations, and environmental data could thus provide valuable insights into diving physiology, stress responses, social interactions, foraging behaviour, and responses to anthropogenic disturbances in free ranging cetaceans. Beyond basic physiology, such approaches will contribute to welfare assessment, veterinary monitoring, and conservation programs for both managed and wild populations.

